# Towards the identification of the molecular toolkit involved in scale worm bioluminescence (Polinoidae, Annelida)

**DOI:** 10.1101/2024.01.28.577566

**Authors:** Carlota Gracia-Sancha, María Conejero, Sergio Taboada, Daniel Martín, Ana Riesgo, Mandë Holford, Aida Verdes

## Abstract

**Background:** Bioluminescence, or the ability of a living organism to produce light, has evolved independently in numerous taxa inhabiting a panoply of ecosystems, although it is more frequent among marine animals. Scale worms are a group of marine polynoid annelids characterized by having dorsal scales, known as elytra, capable of emitting bioluminescent light by a mostly unknown molecular mechanism that may involve a photoprotein called polynoidin. Here, we used RNA-seq data to characterize the expression of genes potentially involved in light production in the polynoid species *Harmothoe imbricata* (Linnaeus, 1767) and *Harmothoe areolata* (Grube, 1860) across tissues of the specimens. We also compared the transcriptomes of the selected species with other bioluminescent and non-bioluminescent polynoids, to identify shared orthologous genes potentially involved in light production. In addition, we investigated the disposition of the photocytes on the elytra using confocal microscopy and histological analyses.

**Results:** Our results showed a total of 16 candidate genes, 15 orthologous genes and 12 enriched GO terms potentially involved in bioluminescence, including genes related with oxidative stress, cytoskeleton, nervous system, stress response, wounding response, eye constituents and metabolic pathways. We also confirmed the presence of photocytes in both species, which appeared distributed around the elytrophore.

**Conclusions:** Among the genes found potentially implicated in bioluminescence we suggest that the oxidoreductase protein, peroxidasin, could be a polynoidin candidate since it appears overexpressed in the elytra of both species and it is located in the endoplasmic reticulum, where this photoprotein has been described to be found.

## Background

Bioluminescence, the ability to produce light by a living organism, is widespread through the Tree of Life. The production of bioluminescence normally involves the oxidation of a light emitting photoprotein, that in some organisms is called luciferin, in a chemical reaction catalysed by an enzyme, called luciferase, resulting in the emission of a photon [1, 2]. Biological mechanisms behind light emission have evolved independently in both terrestrial and aquatic organisms from polar to tropical regions and from coastal waters to the open ocean [3], being more frequent among marine dwellers [4]. There are more than 10,000 species that have disparate mechanisms for the production of light [5], the majority of which are still unknown. The ability to produce light has evolved independently in the different organism lineages, resulting unique mechanisms through which the different taxa produce light. This has led to diversity not only in how each taxon produces light but also in the wavelengths emitted and the specific functions associated with light emission [4–11]. Although there is diversity in the chemical nature of the light-emitting proteins responsible for the production of light, the generic terms of luciferase for the enzymes involved and luciferin for the substrate have become widely accepted [3]. The most studied luciferin was originally described from fireflies, and since then only a few have been characterized, some of which are shared by different lineages [1, 12, 13]. The chemical reaction involved in bioluminescence should be sufficiently energetic to produce an excited molecule that will generate a visible photon when it relaxes, and oxidation reactions fit well into this description, with the breakdown of peroxide bonds among the most widespread mechanism [3]. In this sense, most animal lineages possess different kinds of luciferases for light production [5].

Only in the phylum Annelida, there are more than 100 bioluminescent species described to date [11]. Among these, there are important differences in the chemistry of the bioluminescence systems used by marine and terrestrial taxa, which suggests the independent origin of bioluminescence [14–18]. Polynoids, also known as scale worms, are a group of marine annelids with several bioluminescent species reported, including members of the genera *Gattyana*, *Malmgrenia*, *Polynoe* and *Harmothoe* [11, 16, 19–21] and possibly *Neopolynoe* [10]. Polynoids are characterized by a series of paired structures denominated scales or elytra which are arranged in two rows covering entirely or partially the length of the dorsum [22] (Fig. 1). The elytra are dorsal cirri modified into flat, circular disc-like plates that are attached to the body by a thin layered structure called elytrophore (Fig. 1). The scales serve a wide variety of functions, including egg brooding, facilitating water circulation and also enabling defensive behaviors [23, 24]. When threatened, luminous polynoids emit flashes of light to startle predators that come from the ventral epithelium of the elytra, which has a layer of bioluminescent cells called photocytes not found in non-luminous species [19, 25]. If the stimulus is strong, one or more glowing elytra might be detached from the insertion zone of the elytrophore while the animal swims away [16, 19, 24–26]. The elytra, despite not being attached to the body anymore, continue producing flashes of light for some time, catching the attention of the predator that follows the light source, allowing the polynoid time to escape [24]. Under extreme danger, polynoids might autotomize the entire posterior end of the body. When this happens, the elytra of the posterior end emits light that attracts the predator while the rest of the animal remains dark and escapes to later regenerate the missing segments [19, 27]. Therefore, bioluminescence in scale worms is thought to serve mainly as a warning or distracting mechanism for defense, as elytra are easily autotomized and emit bright flashes of light for some time, acting as a sacrificial lure and allowing the animal to escape avoiding potential threats [1, 16, 19, 24, 28].

**Figure 1.**
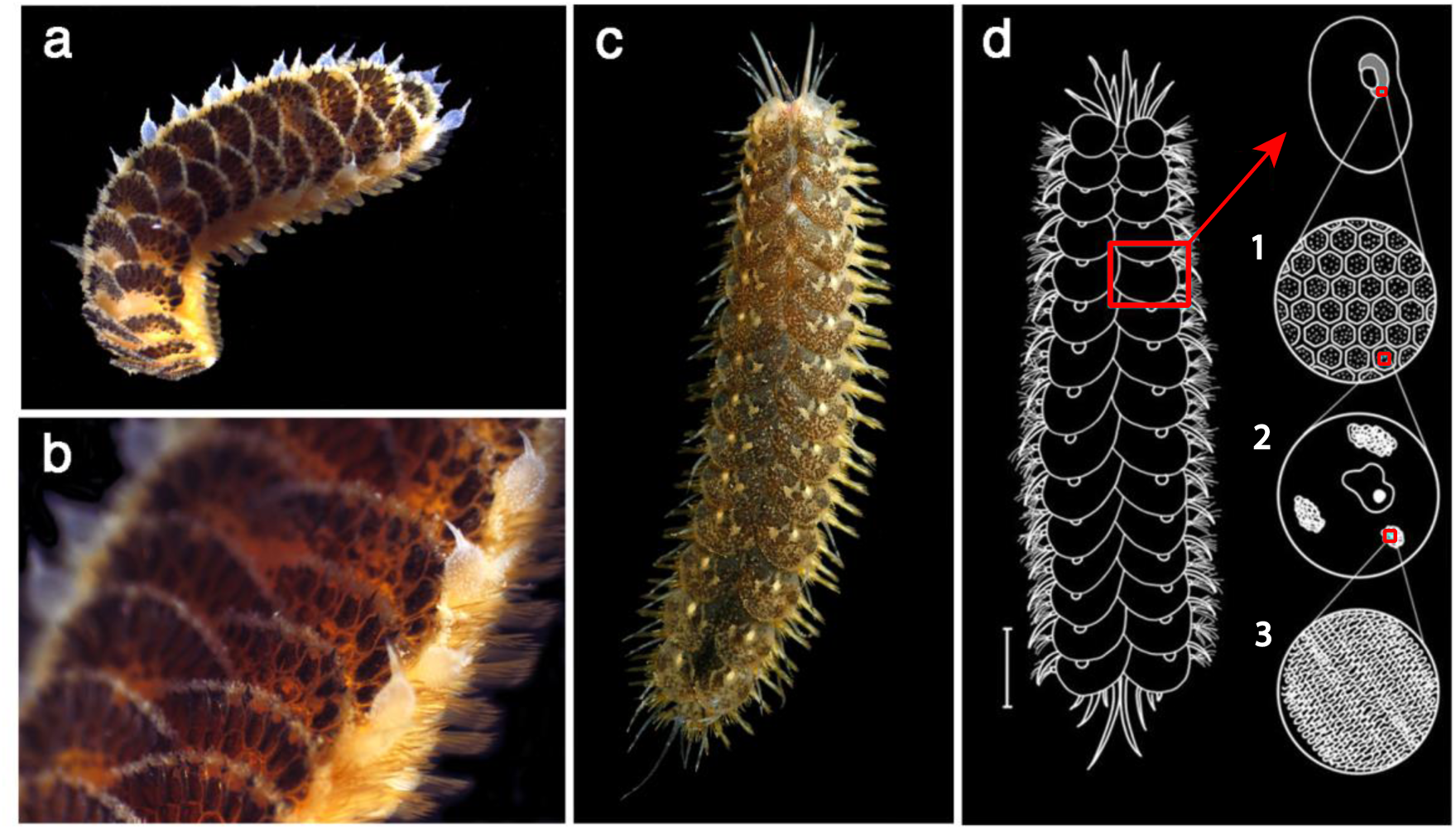
Photographs of *H. areolata* and *H. imbricata* with a diagram representing the bioluminescence system. (**a-b**). Images of *Harmothoe areolata* with a close-up of the dorsum with the elytra in detail. (**c**). Dorsal view of *Harmothoe imbricata*. (**d**). Diagram representing an individual of the polynoidae family showing the disposition of the bioluminescent system. The shaded zone of the elytra corresponds to the bioluminescent area. (**1**). Zoom of this area where the photocytes are arranged in a mosaic pattern. (**2**). A magnified photocyte with three photosomes and a nucleus. (**3**). The paracrystalline structure of the endoplasmic reticulum of the photosomes. (Scale = 5 mm). Modified from *Ouldali et al., 2018* [31].

Morphologically, elytra are composed of a unicellular epidermis, which is interrupted by the insertion of a muscular elytrophore, covered by a cuticular layer. A main nerve trunk traverses the elytrophore branches into nerves that cover the whole structure. The efferent nerve fibres from the main nerve trunk are connect to the photogenic cells, known as photocytes [19] located ventrally, around the elytrophore zone. Each photocyte contains 30 to 50 fluorescent photosomes, the bioluminescent organelles, arranged around the nucleus of the photocytes [29] and linked by various junctions [30]. The polygonal-shaped photocytes exhibit a granular structure composed of undulating microtubules (Fig. 1d), considered to produce secretions implicated in bioluminescence production [29].

Light production in scale worms involves a membrane photoprotein of about 65 kDa called polynoidin, located in the membrane of the photosomes [32] and triggered by superoxide or hydroxyl radicals in the presence of Ca^2+^ and Fe^2+^[30, 32]. The oxidation of the reduced flavin present in the photosomes is hypothesized to produce the superoxide or hydroxyl radicals [16, 29] to trigger light production. Interestingly, polynoidin is also present in non-luminescent scale worms suggesting that bioluminescence might have originated from a mechanism able to quench superoxide radicals [25]. Previous studies of polynoid bioluminescence have been mostly focused on the morphology and ultrastructure of the elytra, the biochemistry of light emission by polynoidin, and the electrophysiology of the photogenic epithelium [19, 25, 30, 32–35]. However, the genetic machinery behind bioluminescence in this group of marine annelids is currently unknown. Here, we take advantage of RNA-Seq based differential gene expression analysis to identify genes potentially involved in scale worm bioluminescence and investigate the morphology of the elytra and the structure of the bioluminescent system in two polynoid species: *Harmothoe imbricata* (Linnaeus, 1767) [36] and *Harmothoe areolata* (Grube, 1860) [37]. For this purpose, we generated RNA-Seq libraries of both elytra and the posterior end of the body and performed differential expression analyses to identify candidate genes potentially involved in light production. Our results suggest that a potential candidate for polynoidin could be a peroxidasin homolog that has been found overexpressed in the elytra of both studied bioluminescent species. Furthermore, numerous transcripts were identified in connection with secondary aspects of the bioluminescent mechanism, encompassing those linked to a response to injury and stress, as well as others correlated with the composition of the photosomes.

## Results

### Transcriptomic and differential expression analyses

The sequencing of the six libraries prepared for *Harmothoe imbricata* resulted in a total of 96.9 million raw reads. The filtered reads examined with FastQC were assembled *de novo* producing a reference transcriptome of 384,174 transcripts including 303,091,081 assembled base pairs (bp) and an N50 value of 1,439 [see Additional file 1]. The sequencing of the six libraries prepared for *H. areolata* resulted in a total of 153.4 million raw reads. The filtered reads examined with FastQC were assembled *de novo* producing a reference transcriptome of 513,553 transcripts including 377,970,784 assembled base pairs (bp) and a N50 value of 807 [see Additional file 1]. The completeness of the transcriptomes based on BUSCO values was high compared to that of other transcriptomic analyses of annelids [38].

The differential gene expression (DGE) analysis for *H. imbricata* resulted in a total of 61,785 transcripts differentially expressed, of which 331 were upregulated in the elytra (Fig. 2a). Of the upregulated transcripts, only 111 could be functionally annotated [see Additional file 2]. To identify genes potentially involved in bioluminescence production, we focused our downstream analyses only on the annotated transcripts differentially expressed in the elytra. The DGE analysis for *H. areolata* resulted in a total of 72,161 transcripts differentially expressed [see Additional file 2], of which 475 were upregulated in the elytra (Fig. 2b). Of the upregulated transcripts, only 65 could be functionally annotated, and again, we focused on those to identify transcripts potentially involved in bioluminescence.

**Figure 2.**
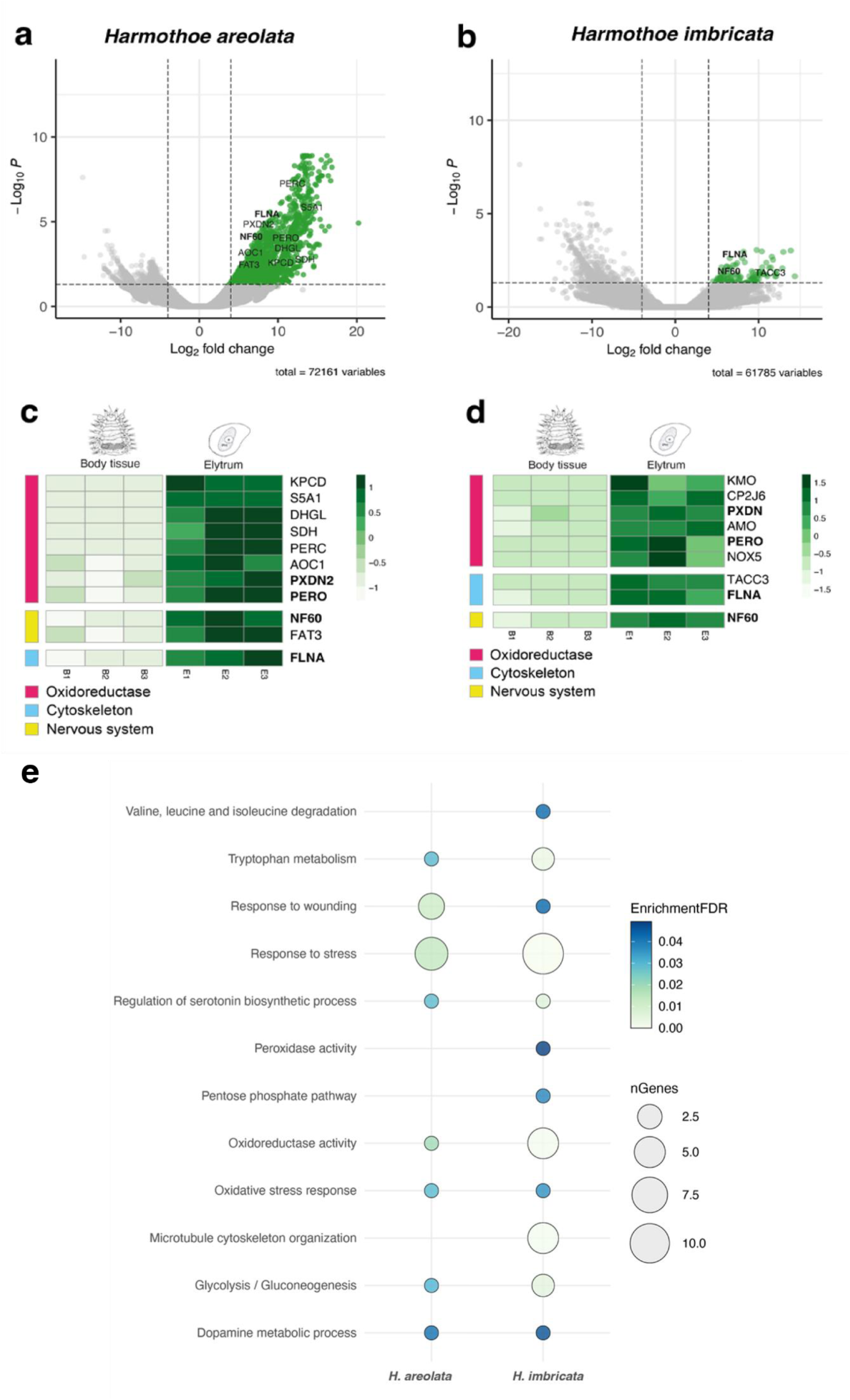
Differential expression and gene enrichment in the elytra of *Harmothoe areolata* and *H. imbricata*. (**a–b)**. Volcano plots displaying the –log10 p value (false discover rate [FDR]) as a function of fold change in the scale and the body of *H. areolata* (**a**) and in *H. imbricata* (**b**). Grey dots in the left represent upregulated genes in the body while green dots at the right correspond to upregulated genes in the elytrum. (**c–d)**. Heatmaps showing selected upregulated genes in the elytrum with their associated functions in *H. areolata* (**c**) and in *H. imbricata* (**d**). (**e**). Gene Ontology enrichment analysis of the upregulated annotated genes in the elytra of both species.

Of the 331 transcripts upregulated in the elytra of *H. imbricata*, only 3 were identified as potentially involved in the bioluminescence production. The annotation of two of these transcripts was related with the cytoskeleton: TACC3 (transforming acidic coiled-coil-containing protein 3) [39] and FLNA (filamin-A) [40]. The remaining transcript blasted against a gene with functions associated with the nervous system, NF60 (neurofilament protein)[41]. In *H. areolata* 475 transcripts were upregulated in the elytra and we selected 11 as possible candidates involved in the bioluminescence process. Several of these transcripts had annotations related to oxidoreductase function: AOC1(amiloride-sensitive amine oxidase) [42], DHGL (Glucose dehydrogenase) [43], PERC (Chorion peroxidase)[44], PERO (Peroxidase) [45], PXDN2 (Peroxidasin homolog) [46], S5A1 (3-oxo-5-alpha-steroid 4-dehydrogenase) [47] and SDH (L-sorbose 1-dehydrogenase) [48]. Another transcript was related with oxygen radical production, KPCD (Protein kinase C delta type) [49]. We also found transcripts annotated to functions in the nervous system, NF60 (neurofilament protein), shared by both species, and FAT3 (Protocadherin Fat 3) [50]. Finally, we found a cytoskeleton related gene, FLNA, which also appeared upregulated in *H. imbricata*.

Heat maps in Figure 2 (c–d) illustrate the expression levels of the transcripts selected in the elytra and the body of both species. We found small differences in the expression profiles of the replicates of each tissue but large differences between tissues, with higher levels of expression in the elytra. For *imbricata*, 6 transcripts were additionally included, given their high expression in the elytra (although they were not differentially expressed). These additional transcripts were annotated to oxidoreductase function: AMO (Putative amine oxidase) [51], KMO (Kynurenine 3-monooxygenase) [52], NOX5 (NADPH oxidase 5) [53] and CP2J6 (Cytochrome P450) [54]. Inside this group we include PERO and PXDN which are also found upregulated in the elytra of *areolata*.

Besides oxidation-reduction processes, in our analyses, we retrieved functional information from 1,149 transcripts of those found upregulated in the elytra in *H. imbricata* and 594 in *H. areolata*, which allowed us to identify various enriched GO terms potentially relevant for bioluminescence (Fig. 2e). The shared pathways with a possible relation with bioluminescence between the two species were: tryptophan metabolism, response to wounding, response to stress, regulation of serotonin biosynthetic process, oxidoreductase activity, oxidative stress response, glycolisys/gluconeogenesis and dopamine metabolic process (Fig. 2e). For *H. imbricata* we found four additional pathways that could be involved in bioluminescence: valine, leucine and isoleucine degradation, peroxidase activity, pentose phosphate pathway and microtubule cytoskeleton organization. For more information of the enriched GO terms [see Additional file 3].

### Identification of shared orthologs between bioluminescent species

The reference transcriptomes of the elytra of *H. imbricata* and *H. areolata* were compared with other bioluminescent and non-bioluminescent polynoid species. With this analysis, we identified orthologous genes shared by bioluminescent species, and present in the elytra of the selected species, but not present in non-bioluminescent species. We focused on the clusters of orthologous genes shared only between the bioluminescent species to identify genes potentially involved in bioluminescence.

All the polynoid species included in the analysis shared a total of 2,994 clusters while *H. extenuata, H. imbricata* (GenBank), and the elytra of *H. imbricata* and *H. areolata* shared 400 clusters of orthologous genes [see Additional file 4]. Of those 400 shared clusters, we identified 15 that could be potentially related to bioluminescence and are listed in Table 1. Interestingly, the Orthovenn analyses revealed a total of 7 clusters related with an oxidoreductase function shared among the bioluminescent polynoids (i.e. Amine metabolic process, hydroquinone, 3-hydroxybutyrate dehydrogenase activity, positive regulation of lipid biosynthetic process, l-ascorbic acid biosynthetic process, thioredoxin-disulfide reductase activity and octopamine biosynthetic process). One cluster exhibited a connection to microtubule organization (i.e. Actin binding), while two clusters were linked to the nervous system, sharing the same GO annotation (i.e. Synaptic transmission, cholinergic), and another two were associated with response to oxidative stress (i.e. Sulfur amino acid metabolic process and regulation of heme biosynthetic process). Furthermore, three clusters demonstrated similarities to genes involved in visual sense (i.e. Estructural constituent of eye lens, retinol metabolic process and visual perception).

**Table 1:**
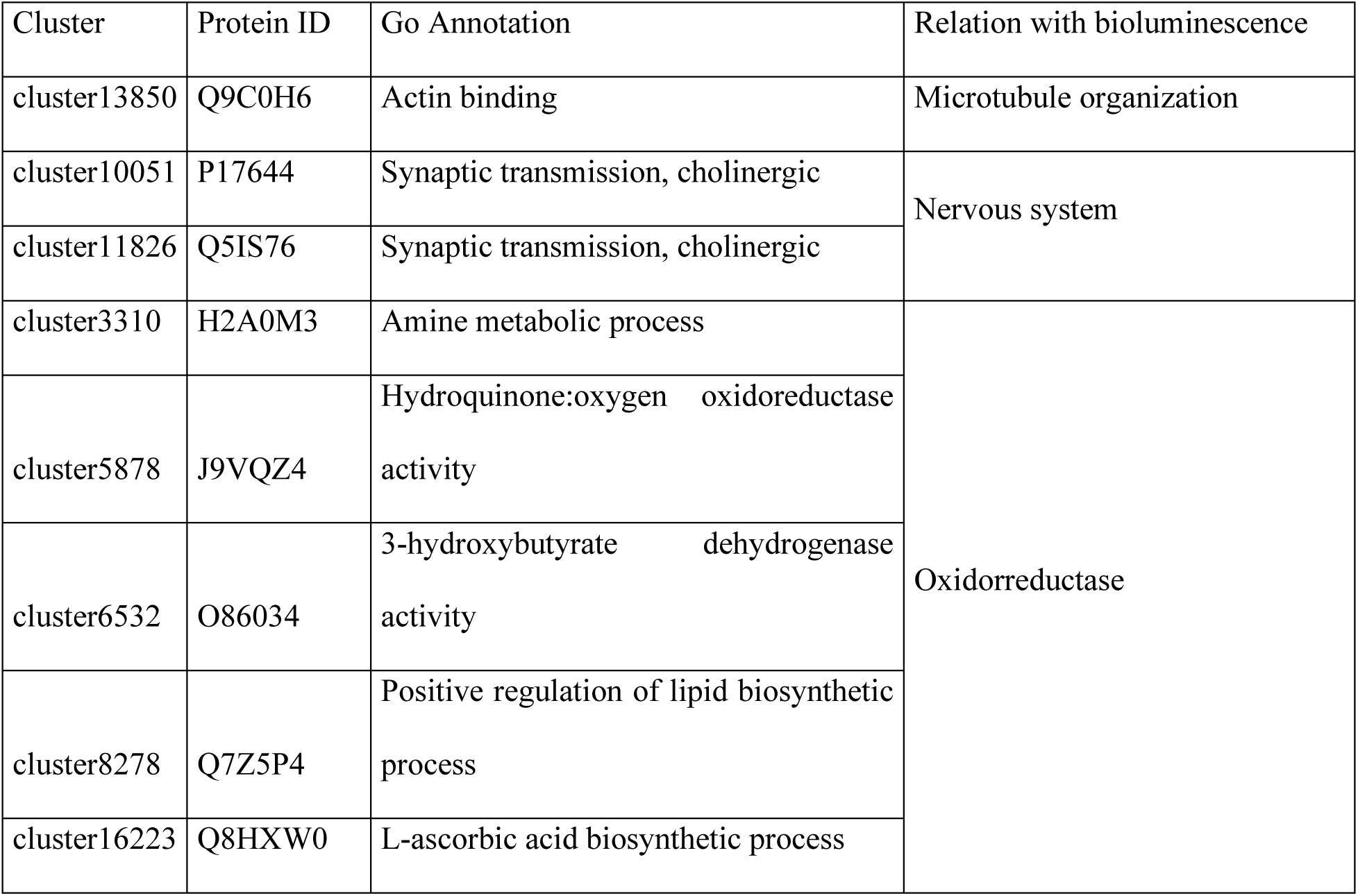
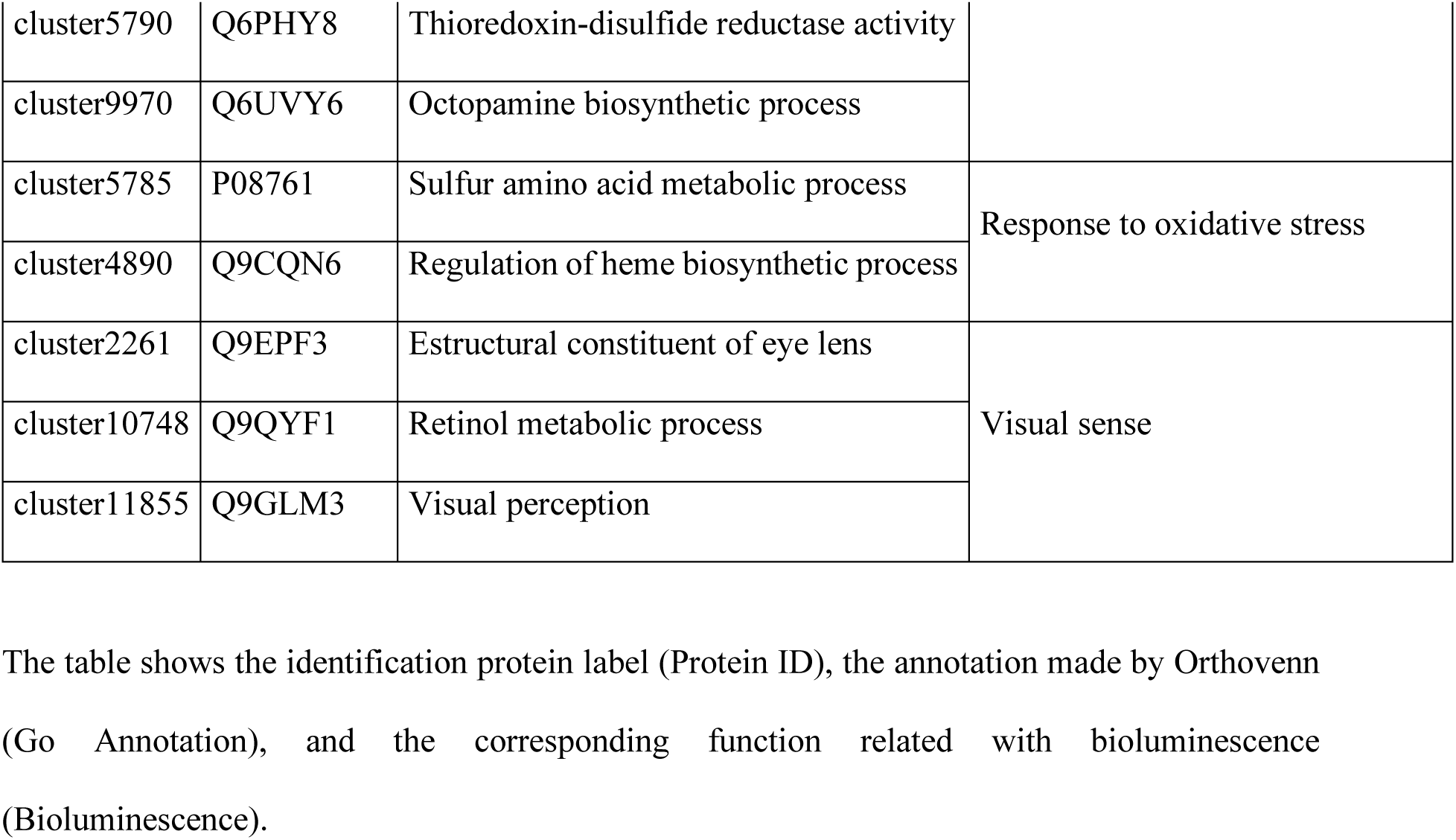
List of clusters shared between the bioluminescent species.

### Histology and tissue organization of elytra

The excitation light used to image the fluorescent photocytes was 405 nm and the fluorescent structures of the elytra emitted within a range spectrum of 517 nm and 570 nm. Figure 3 shows an image of the complete structure of the elytra in *H. imbricata* (Fig. 3a) and *H. areolata* (Fig. 3d) taken from a ventral position. It is known that tubercules covering the surface are autofluorescent, as well as the photocytes [25]. Although the tubercles were located in the dorsal part of the elytra, their autofluorescence was visible from the ventral part. The tubercules covering the dorsum in *H. areolata* were much more conspicuous and with spines, while in *H. imbricata* are rounded (Fig. 3a–f). In figures the ventral side, surrounding the elytrophore we observed a dense mass of brighter fluorescent cells (Fig. 3b–c, e–f). These cells corresponded to the fluorescent organelles of the photocytes, the photosomes, which are distributed within the photogenic area (Fig. 3b–c, e–f). Normally there are 30-50 photosomes per photocyte, with a diameter of 1 - 5 micrometres [29].

**Figure 3.**
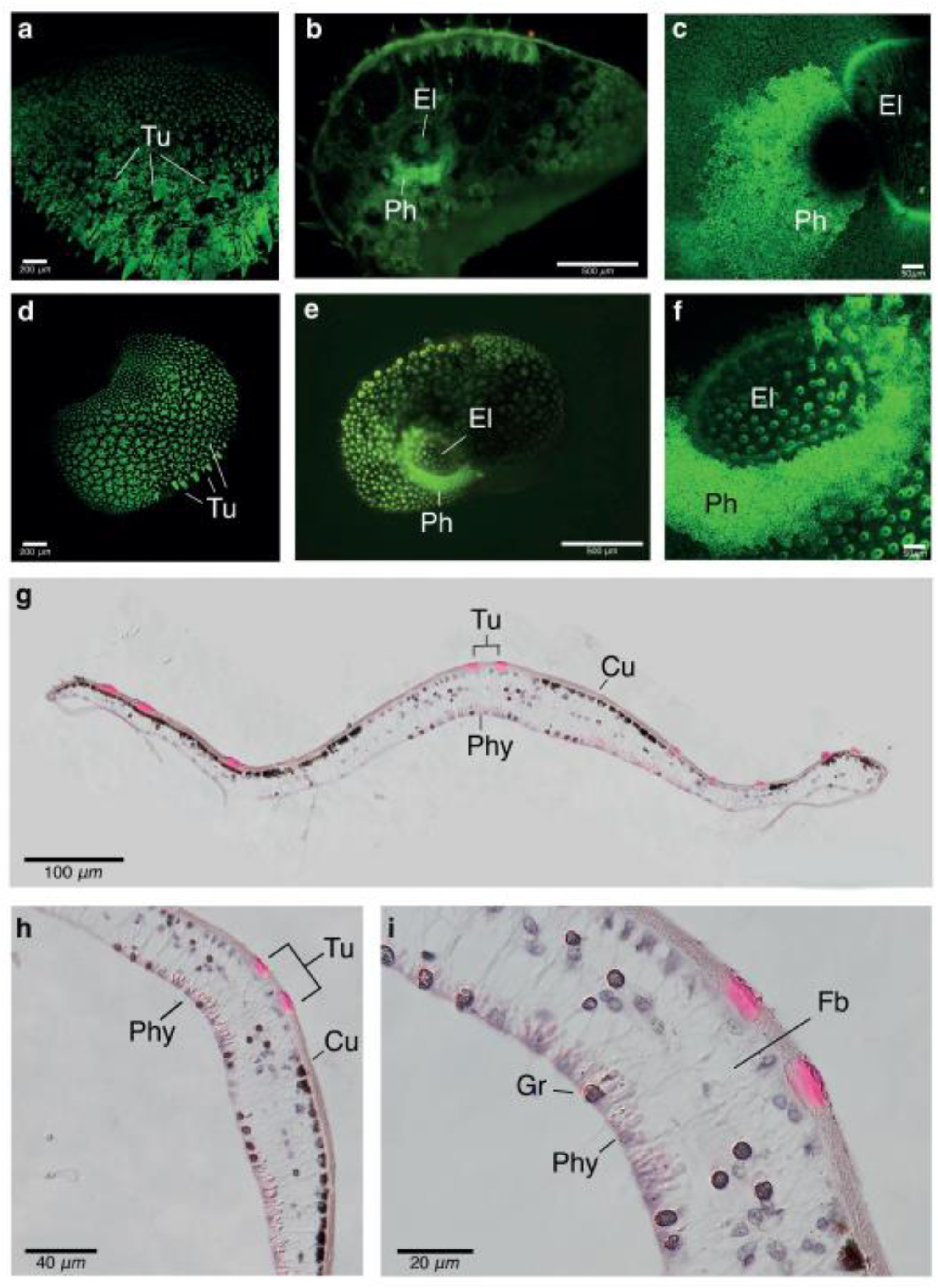
Confocal and light microscopy images of the elytra of *H. imbricata* (a–c) and *H. areolata* (d–f). Absorption spectrum between 517 and 570 nm. (**a**). Dorsal picture of the whole elytrum of *H. areolata* where dorsal tubercules (Tu) are observed as spines. (**b**). Ventral picture of the whole elytrum of *H. areolata* with photosomes (Ph) arranged concentrically around the elytrophore (El). (**c**). Detail of the photogenic area with the photosomes (Ph) in *H. areolata* surrounding the elytrophore (EI). (**d**). Dorsal picture of the whole elytrum of *H. imbricata* where dorsal tubercules (Tu) are observed as small circles. (**e**). Ventral picture of the whole elytrum of *H. imbricata* with photosomes (Ph) arranged concentrically around the elytrophore (El). (**f**). Detail of the photogenic area with the photosomes (Ph) surrounding the elytrophore (El). (**g-i**). Transversal Histological section of an elytrum of H. imbricata stained with haematoxylin-eosin and mounted in DPX. (**g**). Picture of the elytrum the cuticle (Cu) and tubercules (Tu) appearing dorsally and with photocytes (Phy) appearing ventrally. (**h**). Same section at a higher magnification showing photocytes (Phy) at the ventral side breaking their continuity due to the elytrophore. (**i**). Detail of the photocytes (Phy) where the calix shape of these cells is observed. Pigmented granules (Gr) and fibres nerves (Fb).

In the transversal histological sections of the elytra of *H. imbricata* (Fig. 3) we observed several tubercules in the dorsal side (Fig. 3g). Beneath the cuticle we found a continuous upper and lower monostratified epithelium. On the upper layer of the epithelium, we found cell bodies, some of which contained pigment granules (Fig. 3i), nerves (*Fb*) connecting the cuticle with the lower epithelium in a ganglion near the elytral stalk [55], although we could not identify the ganglia in our sections. The lower epithelium was similar to the upper one, with the exception of the presence of photocytes, which were modified epithelial cells, arranged near the elytrophore (Fig. 3i). The insertion of the elytrophore was not clearly identified, although we observed photocytes not following a continuous pattern (Fig. 3h). Photocytes appeared as calyx-shaped cells (Fig. 3i) similar to what has been observed in other species [10].

## Discussion

Our study aimed to identify the genetic machinery involved in the bioluminescence of the polynoid worms *Harmothoe imbricata* and *Harmothoe areolata*. The search of candidate genes with the different analyses led us to the identification of various transcripts in both species that could potentially be involved in light production and additional processes associated with bioluminescence. The DGE analysis allowed us to identify genes that were upregulated in the elytra, the structures where the photogenic cells are located, and thus may be related to bioluminescence production. Additionally, our comparative transcriptomic analysis allowed us to identify orthologous genes shared only by bioluminescent species, which might also be involved in light production. Finally, the morphological analyses with light and confocal microscopy revealed the location and structure of the bioluminescent system within the elytra of both polynoid species.

### Genes potentially involved in the bioluminescence chemical reaction

Bioluminescence in polynoid worms takes place in the elytra. More specifically, this process happens in the photosomes of the bioluminescent cells present in the photogenic area, thanks to the photoprotein polynoidin which is activated by the emission of superoxide or hydroxyl radicals, Ca2+ and Fe2+ [30, 32]. We identified several candidate transcripts and pathways involved in bioluminescence in *H. imbricata* and *H. areolata*, sharing a total of 9 pathways between the two species (Fig. 2e). A total of 11 genes were found differentially expressed and upregulated in the elytra of both species involved in an **oxidoreductase** function. These were identified as the most plausible candidates for luciferase function, given that the primary mechanism operating in bioluminescence involves the breakdown of a peroxide bond [11]. In this sense, one of the most accepted theories of the origin of bioluminescence is the antioxidative hypothesis, and it refers to the evolutionary history of the bioluminescent substrate. Since luciferin has antioxidative properties and high reactivity with reactive oxygen species (ROS), the antioxidative hypothesis suggests that originally, the luciferin was a molecule implicated in reducing oxidative stress [56]. In marine ecosystems, organisms living in shallow waters attempting to escape from predators, might have migrated to the deep ocean where oxidative stress is reduced, leading to a functional change from the antioxidative to the chemiluminescent properties of this molecule [56]. These proteins involved in oxidoreductase processes included Putative Amine Oxidase, Kynurenine 3-monooxygenase, NADPH Oxidase 5 and Cytochrome P450 for *H. imbricata*, while for *H. areolata* these were Amiloride-sensitive Amine Oxidase, Glucose dehydrogenase, Chorion peroxidase, 3-oxo-5-alpha-steroid 4-dehydrogenase, L-sorbose 1-dehydrogenase and Protein Kinase C (Table 2, Additional file 2). Interestingly, both species shared two genes, a peroxidase and a peroxidasin homolog. Oxidoreductase enzymes are involved in controlling the amount of oxygen radicals or reduced compounds that appear in the cells [57]. It is known that the bioluminescent reaction in polynoids is triggered by superoxide radicals that result from the oxidation of riboflavin [16]. Many of the genes selected have NAPH oxidase/dehydrogenase function which are considered one of the major sources of reactive oxygen species [53]. Oxidoreductases are also involved in many other bioluminescent systems including fireflies [58], bacteria [59], fungi [60] and other annelids [61]. Oxygen has been proved to be essential for the luminescent reaction in polynoids [32], therefore, we hypothesize that the oxidoreductase homologs identified here represent candidate polynoidins, the protein responsible of catalyzing the bioluminescent reaction in polynoids. The two only genes that appeared shared by both species are peroxidase (PERO) and peroxidasin (PXDN). The appearance of these genes in the two bioluminescent species studied reinforces the possibility that one of them may be the elusive polynoidin. Moreover, given that the peroxidasin homolog shared is found in endoplasmic reticulum, where polynoidin has been described to be found, this gene appears as the most possible candidate to catalyze the light production in *Harmothoe imbricata* and *H. areolata* [32].

**Table 2.**
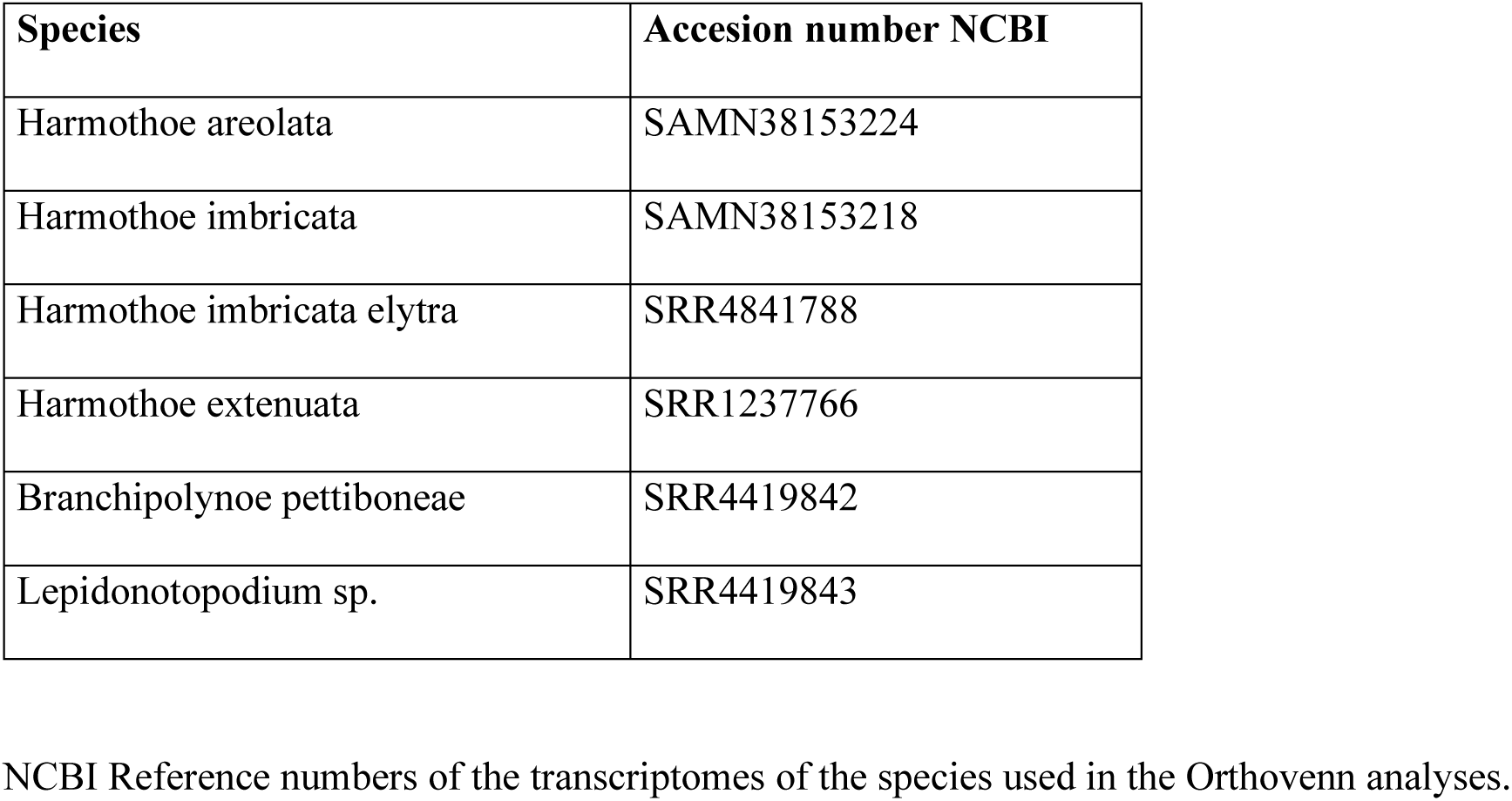
Acession numbers.

Interestingly, two of the orthologous clusters shared by bioluminescent species and one of the upregulated genes in the elytra of both species (NADPH oxidase 5) are involved in **oxidative stress responses**. Oxidative stress is generated by reactive oxygen species [62], between others, which could finally produce bioluminescence in polynoids. It is is possible that initially, bioluminescence occurred as an antioxidant mechanism in order to avoid oxidative stress [56]. This response appearing in the genes overexpressed in the elytra confirms the idea that there are products in the cells of the polynoids capable of acting against this oxidative stress and thus, potentially leading to the production of light.

### Genes potentially involved in additional aspects of the bioluminescence process

Besides the potentially direct production of light of the genes commented above, we identified a battery of genes indirectly related to the bioluminescence process, including those potentially involved in the formation of bioluminescent structures, or with the nervous signal triggering the bioluminescence. Among them, we found two genes differentially expressed in the elytra of the two worms and one shared orthologous cluster associated with the **cytoskeleton.** For both species Filamin-A was upregulated in the elytra. But also, *H. imbricata* showed another gene upregulated in their elytra, Transforming Acidic coiled-coil-containing Protein 3. The cytoskeleton plays an important role in the formation of the photosomes as it is its main structural component. The granules of the photosomes are composed of undulating microtubules that appear regularly arranged in a paracrystalline organization continuous with the endoplasmic reticulum [29]. It has been described that the photocytes of fireflies are also connected to the endoplasmic reticulum, suggesting that this organelle plays a role in the light emission [63, 64]. The task carried out by the gene found homolog to the Transforming Acidic coiled-coil-containing Protein 3 consists in stabilizing the microtubules in the kinetochore fibers of the mitotic spindle [65]. This gene, therefore, could be related with the formation of the paracrystalline organization which conforms the photosomes where the bioluminescent reaction takes part. Filamin-A and the Actin biding cluster, which are shared by both polynoids, are not related with the microtubule organization, but with the actin cytoskeleton. Although the main structural component of the photosomes is indeed the microtubule cytoskeleton, the intermediate filaments of the cytoskeleton, that are formed of actin, are one of the principal elements of the desmosome junctions that join the photosomes inside the photocytes [30]. It is also remarkable that in dinoflagellates, filaments of the actin cytoskeleton are involved in the mechanotransduction of bioluminescence [66]. Consequently, these genes could potentially play an important role in the formation of the photosomes as they have been proved to have relationships with their main structure and with the junctions between them.

One of the most enriched GO categories in both species was **stress response**. The mechanism of bioluminescence in polynoids has hypothesized to be a defensive strategy to escape from predators [67]. It seems obvious that there must be a toolkit to perceive stress and respond creating an impulse ending in light emission to scape. In other studies, similar analyses have also revealed genes overexpressed in the luminal tissue with stress response functions in Terebellidae species [68]. Our study suggests that these genes present upregulated in the elytra of the polynoid worms might work under stress conditions when the animal is trying to escape from its predators. [16][31, 33] We also found several genes related to the **nervous system** that are potentially involved in the bioluminescence process. Bioluminescence in many taxa is controlled by the nervous system, and this includes polynoids. Each elytrum has a central ganglion located near the elytrophore from which nerves branch into nerve fibers in various directions covering the periphery of the elytra [19]. In its upper part, sensory fibers connect with the tubercles or papillae of the cuticle and in its lower part, efferent fibers travel until they reach the photocytes at the base of the elytrum. The stimulus, thus, starts at the upper part and it is transmitted to the central ganglion to reach the photocytes [69]. We identified two upregulated genes in the elytra, Protocadherin Fat 3 and Neurofilament protein and as well as several shared orthologus clusters involved in cholinergic synaptic transmission. Protocadherins homolog, present only in *areolata*, are transmembrane proteins specialized in signaling. Specifically, fat cadherins are defined to enable influence multiple features of neuronal development [50]. This Protocadherin may be involved in the establishment of neuronal connections related to the bioluminescence signal, as these proteins normally binds calcium. Calcium is one of the ions starting the action potential of the luminous process [70] and it could be related to the transmission of the nerve impulse triggering bioluminescence in *H. areolata*. The neurofilament protein, found upregulated in the elytra of both species, is commonly present in neuronal tissue as it is the cytoskeletal constituent of neurons [71]. As it appears overexpressed in both species, this gene may be in fact, one of the genes being responsible of the formation of neurons performing in the elytra. A cholinergic synapse is a chemical synapse that use acetylcholine molecules as neurotransmitter. Nicolas and collaborators in 1978 specified that a cholinergic mechanism was indirectly involved in the luminescence control, and previously it was confirmed a positive acetylcholinesterase histochemical reaction of the elytral ganglion and nerves [72]. All the above mentioned genes may be involved in the neurotransmission necessary for emission of light in polynoids.

Another function indirectly related with the process of bioluminescence is wounding, which its related GO term, **wounding response,** was enriched in the upregulated component of the elytra. When light is being produced, the polynoids detach their luminescent elytra while they are glowing. When this occurs, the luminescent elytra keep on glowing as the same time that the animal tries to scape [24]. The autotomization of the elytra leaves a wound in the worms, that is repaired in the next 15 days, when a new elytrum is grown [73]. Genes capable of responding to this loss of the elytra must be in the elytrum itself, as it must be repaired in order to serve the polychaeta a defensive function.

Three orthologous clusters appeared shared by the bioluminescent species related to **visual perception**. At first it seems strange that genes related to vision appear in the elytra of the polynoids. We must remember that orthologous genes were obtained with the transcriptome of the elytra of our target species and the whole transcriptome of *H. extenuata*. These results suggest that there are homologous genes that are expressed both in the body of the bioluminescent species and in their elytra. With these results we can verify genetically the study carried out by Bassot and Nicolas in 1978 where they confirm that the paracrystalline endoplasmic reticulum characteristic of the photogenic cells in polynoids appears also in the eyes, where it constitutes a greater part of the lens. The relationship that exists in the appearance of the same structures in the photoreceptors as in the photosomes of polinoid species is a mystery to be studied.

Two metabolic pathways related with glycolysis and tryptophan were enriched in the elytra, and even though they have not been reported in bioluminescent systems of polynoids, they have been proved to be involved in other bioluminescent species. The process of obtaining energy out of the glycolysis metabolism, provides energy to an inhibitory mechanism that maintains the photophores in a non-luminescent state in the teleost fish *Porichthys* [74]. In a more closely related species, the earthworm *Lampito mauritii* [75], this metabolic pathway has also been shown to activate a greenish light emission, when the celomic cells where exposed to stimulants and some products of gluconeogenesis [76]. This could indicate that the glucose pathway may interfere in the system of activation and deactivation of light production by polynoids. Also, tryptophan metabolism is central to produce tryptophan compounds, which are one of the main elements of the sea firefly luciferin, cypridinid luciferin [77, 78]. Polynoid’s luciferin has not been discovered yet and since the tryptophan metabolic pathway has been found to be overexpressed in the elytra of polynoids, we propose that tryptophan or its derivatives may be part of the composition of polynoid luciferin.

Finally, we retrieved two more enriched GO categories that were only present in *H. imbricata*: valine, leucine and isoleucine degradation and pentose phosphate metabolism. The generation of NADPH molecules is essential for the bioluminescent process in Polynoidae, as these coenzymes provide reactive oxygen species [16]. Earlier, we indicated the importance of oxidative stress in the reaction of luminescence. Pentose phosphate pathway and valine, leucine and isoleucine degradation are both metabolic pathways in which NADPH is generated [79, 80]. Our study suggests that these pathways may be functioning to generate compounds necessary for beginning the bioluminescent reaction in *H. imbricata*.

### Morphology of the bioluminescent system

Light production in polynoid worms takes place in the photosomes, organelles located in specialized cells called photocytes and found in the elytra. The photosomes are autofluorescent and thus are responsible for the fluorescent properties of elytra, having the ability of emit light under specific excitation wavelengths. In addition, the tubercules also show this autofluorescence. First, it was thought that this property was only activated after the bioluminescent reaction was triggered [81] but it has been proved that this fluorescence is permanently present in these structures [25, 35]. Figure 3 shows images taken from individuals preserved in KINFix after three months, confirming therefore that they effectively do not lose their autofluorescence and that it is present independently of bioluminescence light emission. We confirmed that the excitation wavelength that works best to see the arrangement of bioluminescent structures is UV light, as suggested in previous studies [25, 82]. The green light emission spectrum (517-570 nm) is consistent with that reported for this 20 and other species in previous articles (510-525 nm) [10, 16, 32, 83]. We observed two distinctive green autofluorescent zones: the papilla and tubercules and the area around the insertion zone of the elytrophore. The disposition of the photosomes can be observed more accurately on figure 3c and 3f, where they appear as a dense, tangled mass, which may be due to the paracrystalline form of the endoplasmic reticulum from which they are formed. Our histological sections show the disposition and structure of the photosomes, arranged in the centre of the ventral side of the elytra (Fig. 3g, h, i). They are elongated cells that appear together and close to one another. Each photocyte is connected by efferent fibres to the central ganglion of the elytra, responsible for the transmission of the impulse that triggers bioluminescence. These nerves are visualized as the thin transversal lines connecting both epithelia (Fig. 3i). This innervation can also be seen in non-bioluminescent species, although they lack photosomes [84]. In general, our morphological study is consistent with the organization of light producing organs in other polynoids [10, 19, 25, 35, 82, 85].

Although the light production ability of the polynoid photoprotein system could not be assessed, we identified an array of proteins involved in oxidoreductase processes that point to both luciferase and luciferin proteins. Peroxidasin might correspond to the elusive polynoidin, the photoprotein responsible of catalyzing bioluminescence reaction in polynoids. We have seen that the oxidoreductase most likely to be polynoidin is a **peroxidasin homolog**, since it is overexpressed in the elytra of both species and is found in the endoplasmic reticulum, which is an essential part of the structure of photocytes and where this photoprotein has been described to be found. Functional validation analysis must be done to confirm the role of this protein in the bioluminescent system of *Harmothoe.* It will be necessary to search for this supposed polynoidin in other bioluminescent polynoids to verify its role in the bioluminescence process within this genus.

## Methods

### Sample collection and preservation

Six specimens of *Harmothoe imbricata* and six of *Harmothoe areolata* were collected by SCUBA diving in O’Grove (42.498448, −8.865719, Galicia, Spain; 12 m depth) and near Blanes (41.673245, 2.802646, NW Mediterranean Sea; 10 m depth) respectively, and kept alive in seawater for transport to the laboratory. Taxonomic identification was done following the original description of the species and relevant additional literature [36, 86] under a Zeiss Stemi 2000 stereoscope up to 35x magnification by experts on annelid taxonomy. Before sample preservation, bioluminescence emission was confirmed in a dark room by disturbing the individual with tweezers. For transcriptomic analyses, we preserved the specimens in RNAlater (Life Technologies), during 24h at 4 °C, replaced it once, and stored samples at −80 °C until further processing. For histological analyses, samples were preserved in 2.5% glutaraldehyde in 1M PBS and 0.34M NaCl and KINFix, which allows the sample to be used in examination by histological or immunohistochemical techniques [87].

### RNA extraction, cDNA library preparation, and sequencing

For RNA extraction, dorsal elytra and a portion of the posterior end of the body were dissected from three specimens of each of the two species, totalling three replicates per tissue and per species. Total RNA was extracted from the elytra and from the posterior end of the body with TRIzol (Invitrogen), following the manufacturer’s instructions. Total RNA from each of the six biological replicates of *H. areolata* was sent to New York University Center for Genomics and Systems Biology (New York, USA) for cDNA library preparation and sequencing. A total of twelve cDNA libraries were generated with Illumina TruSeq library prep kit v2, corresponding to the three biological replicates of each tissue type. We verified the quality and integrity of the total RNA and the cDNA library with an Agilent BioAnalyzer (RNA and high sensitivity DNA assay respectively). Libraries were then sequenced using Illumina HiSeq 2500 v4 technology, at 150 base-pairs (bp) paired-end reads.

### De novo transcriptome assembly, annotation, and differential expression analysis

The quality of the raw reads generated from each of the six libraries was evaluated using FastQC v0.11.5 [88] and then, adapter sequences and low-quality bases (phred score < 30) were removed using Trimmomatic [89] Processed reads of all replicates were then pooled together and assembled *de novo* using the software Trinity v2.4.0 ([90, 91] to generate a reference transcriptome. Functional annotation of the reference transcriptomes was performed first by blasting the transcripts with DIAMOND [92] against the UniprotKB database [93, 94] with a cut-off E value of 1e-5. Transcripts with blast hits were further annotated using Gene Ontology (GO) terms with BLAST2GO PRO [95].

To identify the genes significantly upregulated in the elytra and the posterior end of the body, a differential gene expression (DGE) analysis was carried out with the Trinity module [90], which incorporates Bowtie and RSEM [96], to map and estimate transcript abundance, and edgeR [97] to identify differentially expressed transcripts [91] in both species. We later selected only those genes showing a minimum of 2-fold change, and *p*-value cut off for FDR of 0.01. A GO enrichment analysis was performed using the GO annotations of the significant differentially expressed transcripts using gProfiler [98] with a FDR cutoff 0.05.

### Comparative transcriptomics of bioluminescent and non-bioluminescent polynoid species

To assess whether bioluminescent polynoids possess a unique set of proteins shared across the family, we compared the transcriptomes of bioluminescent (including our reference transcriptomes) and non-bioluminescent species using OrthoVenn2 [99]. For this, we assembled the reference transcriptomes of the non-bioluminescent polynoids *Branchipolynoe pettibone* (Miura & Hashimoto, 1991) [100] and *Lepidonotopodium* sp. [100, 101] and the bioluminescent *Harmothoe extenuata* (Grube, 1840) [102], using the available reads in SRA (Table 2) and applying the same bioinformatic pipeline as for our reference transcriptomes. Predicted protein datasets from the transcriptomes were obtained with TransDecoder [90] and implemented in OrthoVenn2 where we set a cut-off E value of 1e-5 for the comparisons. This served us to select only those orthologous gene clusters shared by the luminous polynoid species (“bioluminescent clusters”) for downstream analyses. Functional annotation of the shared bioluminescents clusters was automatically performed in OrthoVenn2 against UniProtKB [93, 94] as well an additional GO enrichment analysis.

### Histology analysis of elytra

To identify the presence of photocytes, histological analyses were performed on the preserved elytra of *H. imbricata.* Briefly, samples were rinsed in distilled water, dehydrated through an ascending series of ethanol, bathed in xylene and embedded in paraffin overnight at 60°C. Serial 7µm-thick sections were then made with a Leitz 1512 microtome and stained with a standard haematoxylin - eosin protocol and mounted with DPX. The resulting preparations were scanned using an Olympus© BX51-P microscope and photographed with an Olympus DP-23 camera.

Confocal and light microscopy of elytra preserved in formalin was also performed taking advantage of the autofluorescence of the photosomes present in the photocytes of the elytra of both species. The excitation light used was of 405 nm in order to observe the disposition and arrangement of the photocytes within the elytra. The confocal microscope used was an LSCM Olympus FV1200 at Universidad Complutense de Madrid and the light microscope was a Olympus BX53 at Museo Nacional de Ciencias Naturales (MNCN).

## Supporting information

Sequencing information. De novo transcriptome assembly metrics and transcriptome completeness statistics based on BUSCO analysis with the Metazoa

Scale up subset. Table showing the results of the Differential Expression Analyses performed for both species (one species per sheet)

GO terms. Results of the Gen Ontology analyses performed with the whole transcriptome of the selected species as background.

Orthovenn results. Orthologous genes shared by the selected bioluminescent species.

## Acknowledgements

We acknowledge Ana Sánchez and Eduardo Roldán for help with fluorescent microscopy and the staff of the imaging center of Universidad Complutense de Madrid for help with the confocal microscope. We are also thankful to Belén Arias for help amplifying the cytochrome c oxidase I of *Harmothoe* from Galicia to confirm IDs. This work was funded by the Camille and Henry Dreyfus Teacher-Scholar Award and National Institutes of Health (NIH-NIMHD grant 8-G-12-MD007599) to MH, and the European Union’s Horizon 2020 Research and Innovation program through a Marie Sklodowska-Curie Individual Fellowship (grant agreement 841576) and the Spanish Ministry of Science MCIN/AEI/ 10.13039/501100011033, European Union NextGenerationEU/PRTR (grant IJC2020-045256-I) to AV.

## Ethics declarations

### Ethics approval and consent to participate

Not applicable.

### Consent for publication

Not applicable.

## Availability of data and materials

The datasets generated and/or analyzed during the current study are available in the [Additional files] repository, [https://www.dropbox.com/sh/rqx5l06cglk7ojp/AAAdKFH_i9xSo_2_yeYJNpFQa?dl=0]

## Additional files

**Additional file 1 Sequencing information.** *De novo* transcriptome assembly metrics and transcriptome completeness statistics based on BUSCO analysis with the Metazoa gene set.

**Additional file 2 Scale up subset.** Table showing the results of the Differential Expression Analyses performed for both species (one species per sheet).

**Additional file 3 GO terms.** Results of the Gen Ontology analyses performed with the whole transcriptome of the selected species as background. The results for each species are found on separate sheets.

**Additional file 4 Orthovenn results.** Orthologous genes shared by the selected bioluminescent species.

